# Programmed trade-offs in protein folding networks

**DOI:** 10.1101/2020.04.07.030056

**Authors:** Sebastian Pechmann

## Abstract

Maintaining protein homeostasis, i.e. a folded and functional proteome, depends on the efficient allocation of cellular protein quality control resources. Decline and dysregulation of protein homeostasis are directly associated to conditions of aging and neurodegeneration. Molecular chaperones as specialized protein quality control enzymes form the core of protein homeostasis. However, how chaperones selectively interact with their substrate proteins thus allocate their overall limited capacity remains poorly understood. Here, I present an integrated analysis of sequence and structural determinants that define interactions of the *Saccharomyces cerevisiae* Hsp70 Ssb. Structural homologues that differentially interact with Ssb for *de novo* folding were found to systematically differ in complexity of their folding landscapes, selective use of nonoptimal codons, and presence of short discriminative sequences. All analyzed characteristics contributed to the prediction of Ssb interactions in highly complementary manner, highlighting pervasive trade-offs in chaperone-assisted protein folding landscapes. However, short discriminative sequences were found to contribute by far the strongest signal towards explaining Ssb interactions. This observation suggested that some chaperone interactions may be directly programmed in the amino acid sequences rather than responding to folding challenges, possibly for regulatory advantages.

## Introduction

Correctly folded proteins are required for virtually all biological function. Inside the crowded cellular milieu, the ability of proteins to fold into their functional conformations strongly depends on interactions with their environment (*1–3*), notably with helper proteins called molecular chaperones (*4, 5*). Chaperones as specialized protein quality control enzymes form the core of an elaborate regulatory and quality control system, the protein homeostasis network, that safeguards that all proteins are correctly folded while avoiding aberrant aggregation (*6*). Through direct protein-protein interaction, chaperones assist their client proteins with *de novo* folding, avoiding aggregation, degradation, and translocation (*4, 5*). Failure of protein homeostasis leads to the accumulation of misfolded proteins and is directly associated with conditions of aging (*7–10*) and aging-related neurodegeneration (*11*). Importantly, the observation that an overall limited chaperone capacity within cells has strongly co-evolved with proteome size and complexity (*12,13*) suggests that an efficient use of cellular resources is prerequisite for maintaining a folded proteome, i.e. protein homeostasis. However, the principles of specificity and selectivity in chaperone interactions that govern their efficient allocation remain poorly understood.

Generally, chaperones bind to hydrophobic sequences in aggregation prone and folding challenged proteins (*14–17*) that are normally protected by the native structure. Interactions with posttranslationally acting chaperones such as the ubiquitous heat shock protein Hsp90 are thus tightly coupled to protein thermodynamic stability that limits spontaneous unfolding and exposure of such sequences (*18*). In turn, cotranslationally acting chaperones, in *S. cerevisiae* foremost the ribosome-associated Hsp70 chaperones Ssb1 and Ssb2, directly bind to elongating nascent polypeptides at the ribosome exit tunnel to facilitate their *de novo* folding (*15,17,19–21*). The redundant Ssb1 and Ssb2, or collectively Ssb, interact with largely overlapping sets of proteins (*15, 22*) through recognition of linear peptide sequences of 5-7 amino acids (aa) in unfolded or partially folded polypeptides (*16,17*). Remarkably, in long multi-domain proteins, these binding events directly match the domain structure of the substrate proteins thus promoting their folding (*23*). In addition, interactions of cotranslationally acting chaperones can be modulated by local translation elongation kinetics (*23, 24*) as well as prelocalization of protein-coding mRNAs to specialized ribosome-chaperone complexes (*25*).

A unique characteristic of chaperones is their remarkable ability to interact with hundreds of client proteins yet maintain selectivity (*15,18, 26, 27*). Normally, such broad specificity in molecular recognition is achieved through plasticity in the receptor protein. For instance, hubs in protein networks interact through multiple binding sites on their surfaces (*28*). Similarly, a flexible loop in the antibody binding site that can assume multiple conformations permits binding to many different structures (*29*). The highly specialized eukaryotic chaperonin TRiC/CCT equally utilizes multiple binding sites to achieve specificity (*30*). In contrast, Ssb interacts selectively with over a thousand proteins on the ribosome (*15, 22*), presumably through its main binding domain. Understanding the determinants of specificity and selectivity of ribosome-associated chaperones is of particular interest as they are directly coupled to the sequence determinants of *de novo* protein folding in the cell (*21,31*). Importantly, the integration of protein structures into chaperone interaction maps has highlighted that the structural fold of proteins alone is generally not a determining factor of chaperone interaction (*18,26,32*). Thus, through tremendous progress on understanding chaperone action and interaction, it has also become increasingly clear that not a single feature but rather the complex interplay of multiple protein characteristics determines their interaction with chaperones.

To this end, trade-offs are omnipresent in biology and equally constitute an inherent part of the complexity of proteins (*33*). Foundational has been the realization that, in principle, all protein can form aggregates (*34*). Accordingly, it has become clear that the successful folding of proteins that are generally only meta-stable (*35*) and marginally soluble (*36*) always stands in direct competition with unwanted protein aggregation (*37*). This competition is governed by the complexity of the folding landscape, thermodynamics, and folding kinetics: the potential for misfolding through formation of non-native contacts, the thermodynamic stability of the native state, as well as the kinetics of reaching the native state before any misfolding and aggregation events are all critically important (*38–41*). Small and extremely fast-folding proteins can reach their native states within milliseconds without chaperone assistance (*42*). However, the potential for non-native contacts can readily impact folding dynamics (*43*). Moreover, while protein topology strongly influences available folding pathways (*44*), stark differences in folding exist even between structural homologues. Pervasive frustration in protein structures (*45*) suggests that proteins are rarely optimized for structure alone but instead serve to accommodate diverse functional constraints (*33*). Certainly, a fine balance between folding and functionality is required (*46*). For instance, too much protein stability can negatively affect protein solubility (*47*) and function (*48,49*) such as inhibition of enzyme activity through hampering of protein dynamics (*50,51*). Conversely, mutations that lead to functional innovation are on average detrimental to protein stability (*52*). Similarly, specificity in protein interaction (*29*) may constrain the use of specific amino acids for molecular recognition (*53*) or require negative design principles that prevent unwanted interactions (*54*). Thus, multispecificity for several interaction partners usually comes at a cost, for instance to protein stability (*55*). Chaperones are poised to play a pivotal role in balancing trade-offs between protein folding and function. Consequently, the principles of chaperone interaction extend in importance beyond their role in selective protein quality control.

Here, I present an integrated analysis of trade-offs that govern the interactions of *S. cerevisiae* single-domain proteins with the cotranslationally acting Hsp70 chaperone Ssb for de novo folding. Through systematic comparison of structural homologues that differentially interact with Ssb, contributions of the complexity of the folding landscape, presence of discriminative sequences, and codon usage are analyzed. Chaperone interactors were found to possess on average more risky folding landscapes as evident by lower contact density and longer contact distance. Selective use of nonoptimal codons that may promote formation of critical hydrophobic contacts appeared to compensate for chaperone assistance. However, the strongest determinant of Ssb interaction was the presence of discriminative short sequences within proteins. This observation suggested that some chaperone interactions may be directly encoded in the protein sequences rather than reflecting a folding requirement. Finally, the present analysis highlights the importance of an integrated and contextual understanding of chaperone-assisted protein folding wherein some chaperone interactions are directly programmed, likely for functional or regulatory advantages.

## Results

The prevalence of structural homologues that differ in their chaperone dependence clearly indicates that the protein fold alone is in general not a stringent requirement for chaperone interaction (*18, 26, 32*). Comparative analyses of homologous proteins thus allow to delineate any trade-offs within alternate sequences that fold into the same native protein structures. This is especially of interest for understanding the sequence determinants of de novo protein folding and of specificity in chaperone interactions.

### Single domain proteins interact with Ssb in fold-independent manner

Making use of two recent data sets describing the cotranslational interactions of the highly redundant yeast chaperones Ssb1 (*22*) and Ssb2 (*15*), a consensus of Ssb interacting (*I*) and noninteracting (*NI*) proteins was derived (Figure 1A). Both data sets divided interacting proteins into ‘strong’ and ‘weak’ interactors based on their binding enrichment relative to their translation (*15,22*). Here, proteins that strongly, i.e. systematically, interacted with either Ssb1 or Ssb2 were categorized as ‘interactors’ (*I*) while proteins that interacted with neither Ssb, or only weakly, i.e. sporadically, with at most one of Ssb1 or Ssb2 were considered as ‘non-interactors’ (*NI*) (*see Methods*).

**Figure 1:**
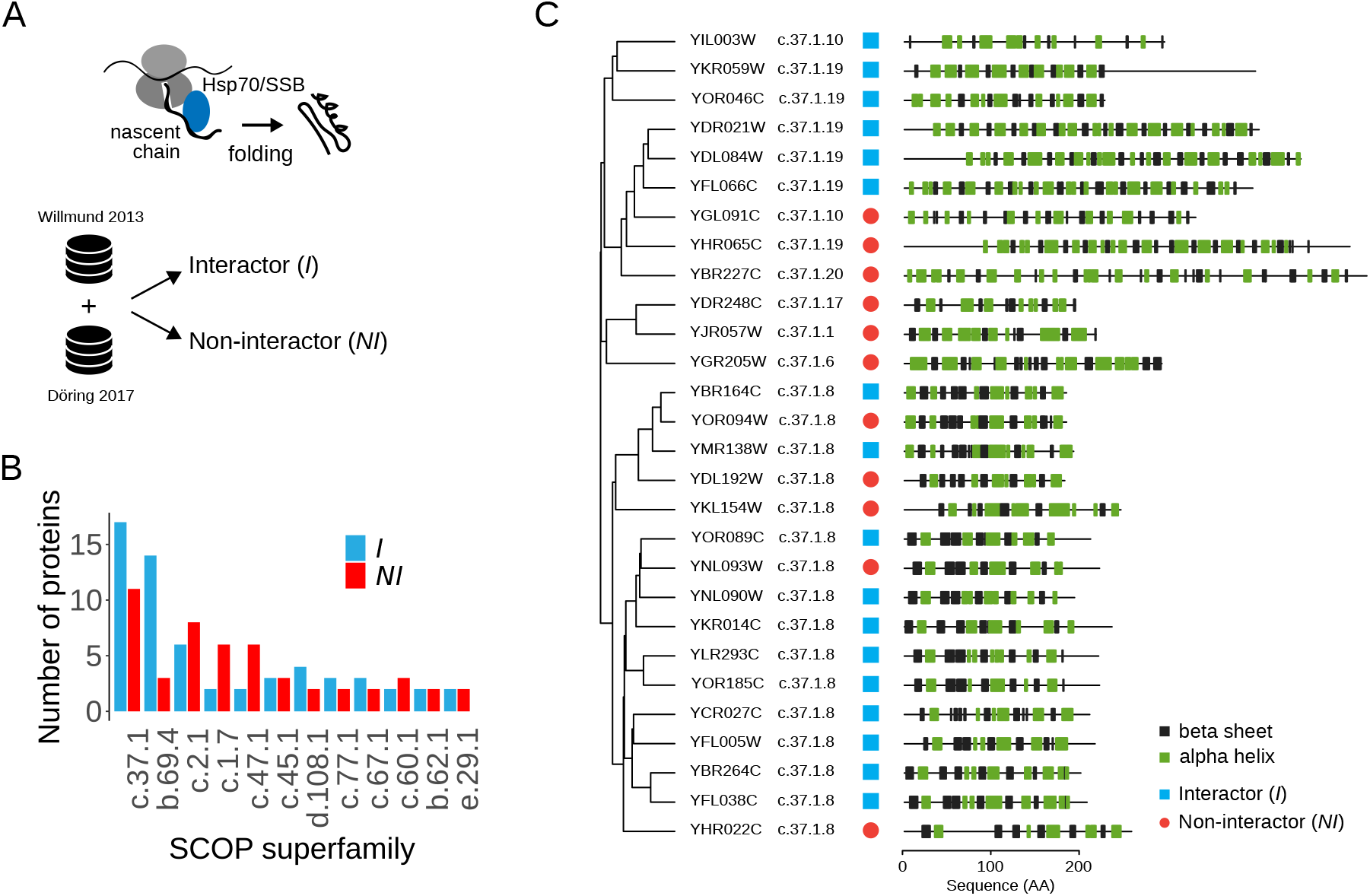
Structural classification of single-domain yeast Ssb1/Ssb2 interactors. **A** Schematic. The ribosome-associated Hsp70 chaperones Ssb1 and SSb2 (Ssb) interact with newly synthesized polypeptide chains to promote their *de novo* folding. Two recent data sets were used to define consensus Ssb interactors (*I*) and non-interactors (*NI*). **B** SCOP superfamily assignment of single-domain *I* and *NI* proteins. **C** Structural phylogeny of the proteins from the superfamily “P-loop containing nucleoside triphosphate hydrolases” (c.37.1). For each protein also indicated are protein family classification, Ssb interaction, and the protein secondary structure profile.

Proteins with multiple domains rapidly increase in complexity. To uncover sequence and structural determinants of Ssb interactors in a setting as controlled as possible, I focused on all single-domain *I* and *NI* proteins that were grouped according to their protein superfamily fold classification (*56*) based on the Structural Classification of Proteins (SCOP) database (*57*) (see Methods). The most populated superfamilies represented abundant protein folds including the ‘P-loop containing nucleoside triphosphate hydrolases’ (c.37.1), ‘WD40 repeat-like’ proteins (b.69.4) and ‘Tyrosine-dependent oxidoreductases’ (c.2.1) (Figure 1B). For all proteins in superfamilies with at least 2*I* and 2 *NI* proteins, available protein structures were retrieved and, for all proteins without experimentally determined protein structures, homology models were built (Table 1, *see Methods*). Within protein superfamilies, proteins subdivide into protein families. A structural phylogeny of the especially well populated c.37.1. fold readily illustrated similarities and differences within this superfamily (Figure 1C, see Methods). Particularly standing out were highly conserved structural similarities within the G-protein family (c.37.1.8). Taken together, the systematic structural classification and homology modeling of single-domain S. cerevisiae proteins prepared for an in-depth comparison of homologous *I* and *NI* proteins to better understand what renders proteins dependent on chaperone assistance for de novo folding.

**Table 1:**
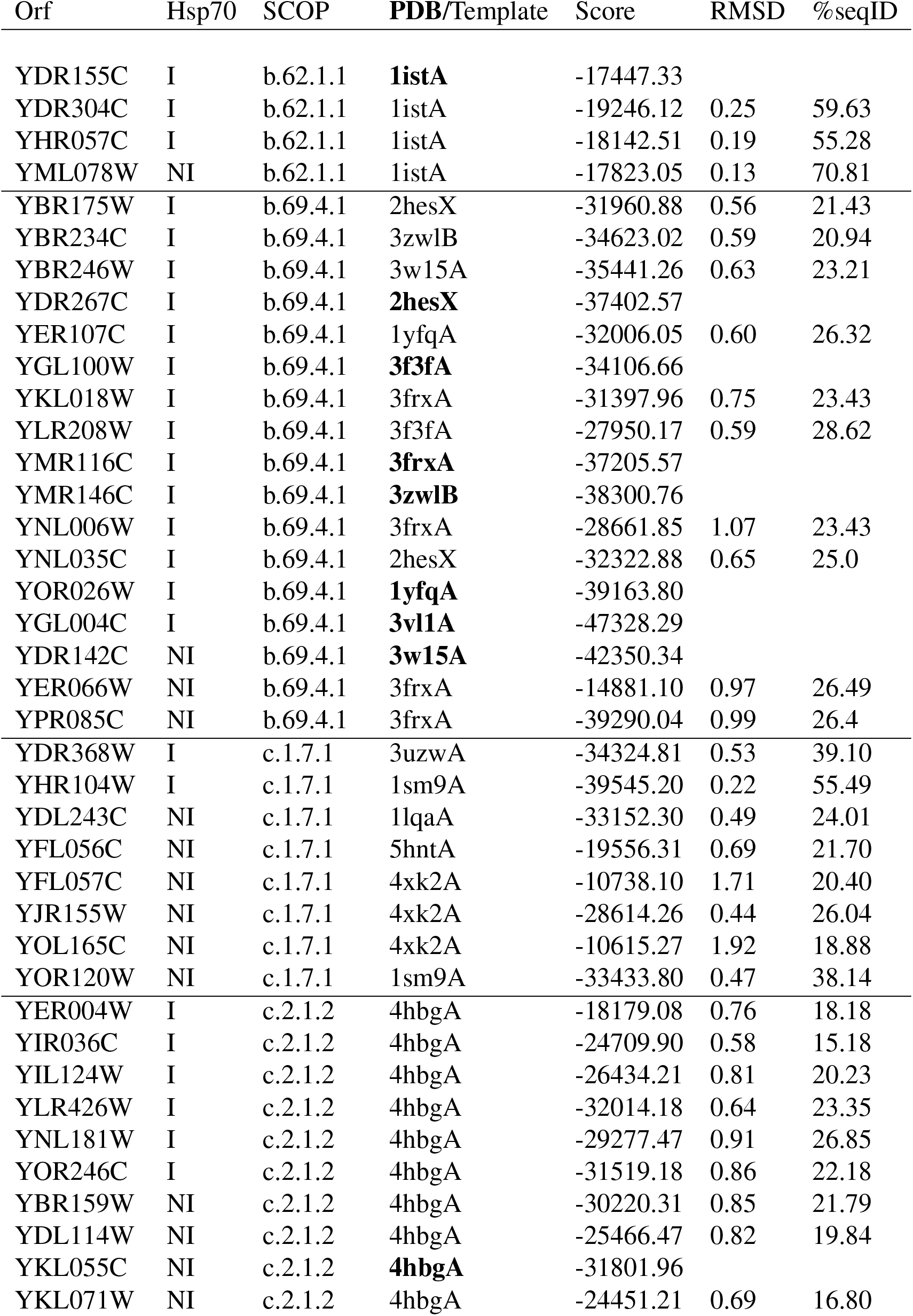
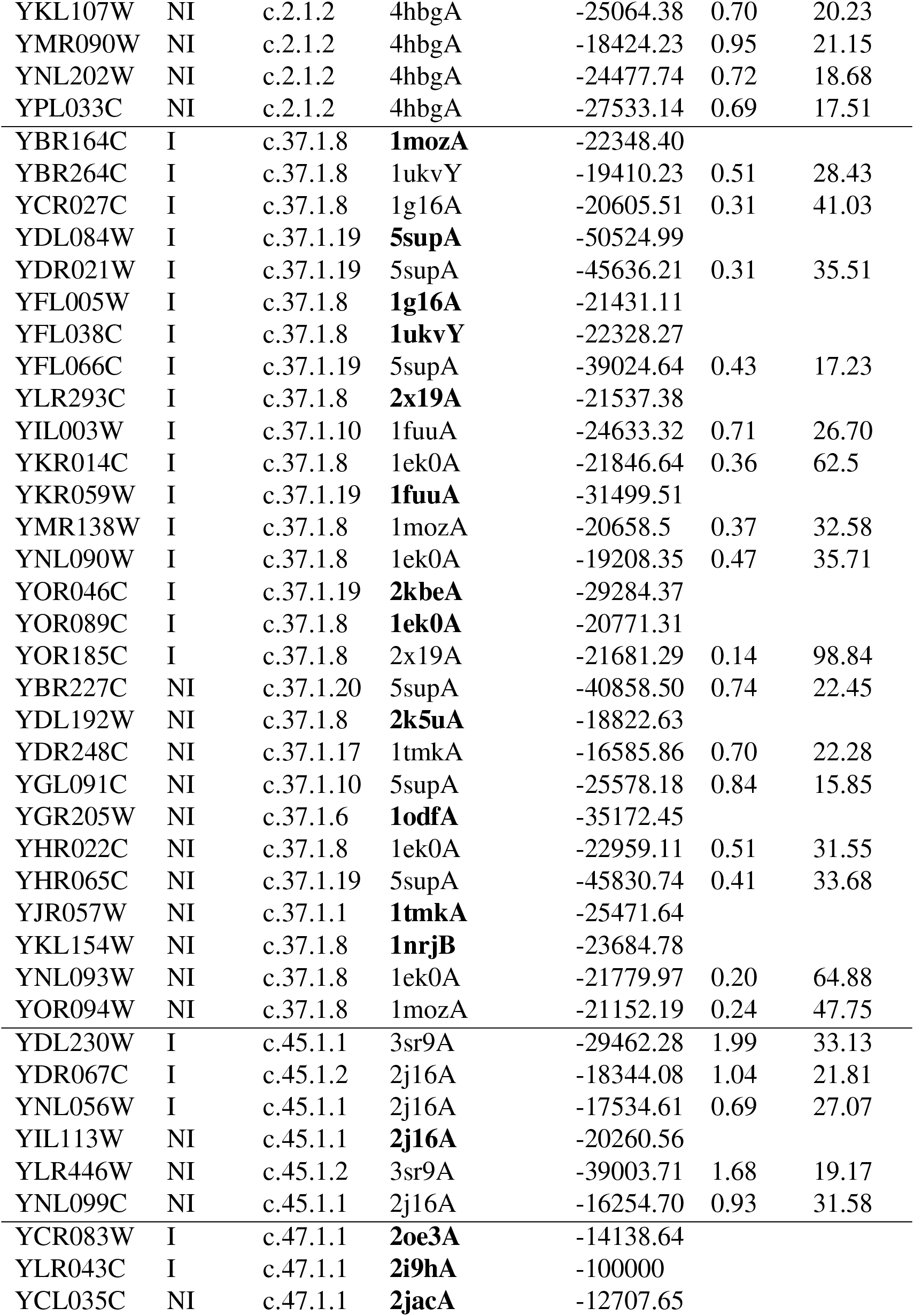
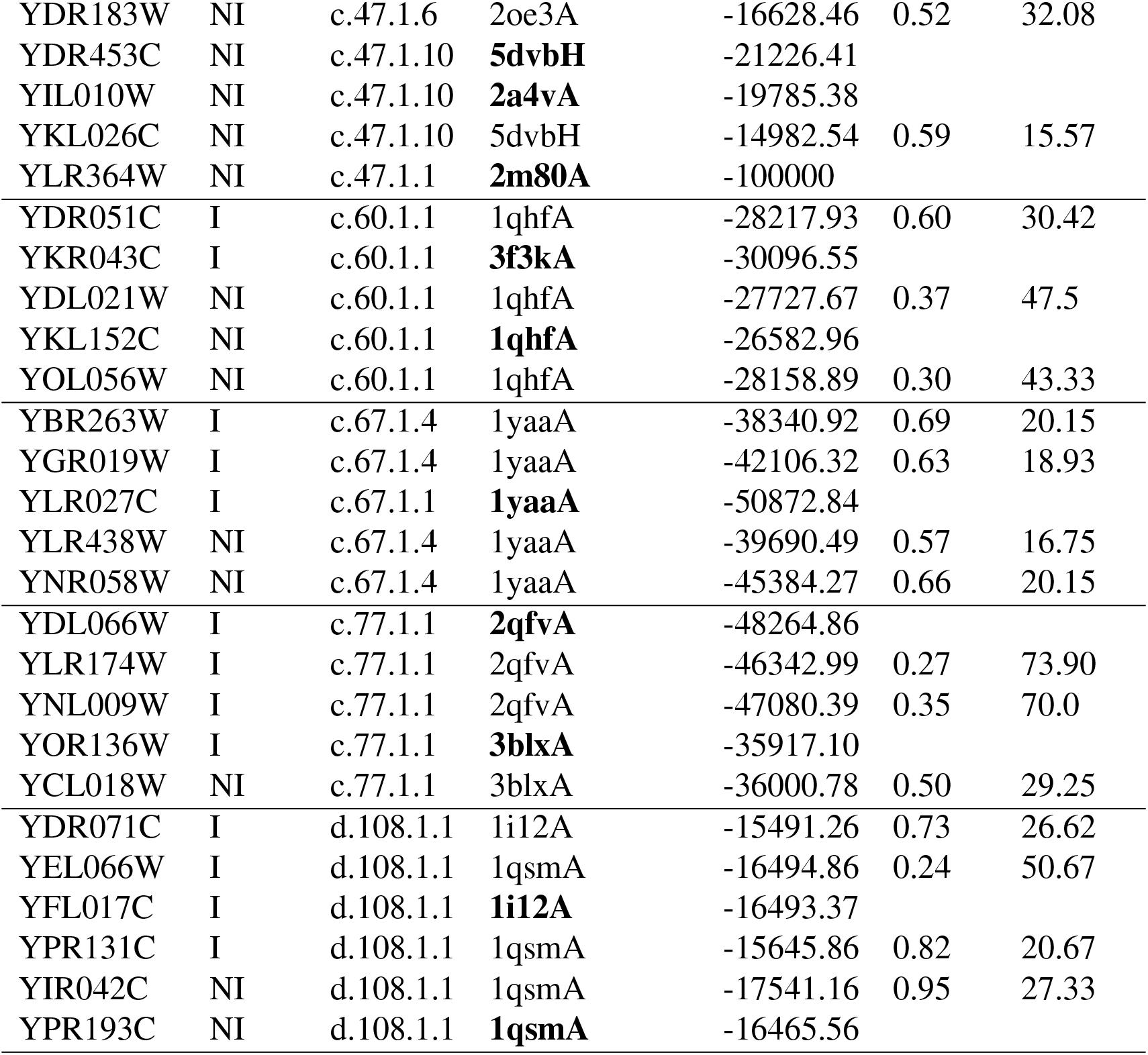
Overview of proteins and protein structures analyzed. All experimentally determined protein structures are listed with their pdb code in bold face and no RMSD or seqID values. The DOPI score resulted from modeling the sequences of proteins with native pdb structures against their own template to add missing residues. All generated homology models are listed with the pdb code of the template, DOPI score of the best scoring model, as well as RMSD and seqID relative to the template.

### No conservation of hydrophobic contacts in structural homologues

I initially focused on the most controlled data at the protein family level, namely a comparison of the 11 *I* and 5 *NI* proteins from the G-protein family (c.37.1.8). Superposition of the 16 protein structures clearly revealed a strongly conserved structural core and overall very high structural similarity (Figure 2A). This observation quantified to very low pairwise root mean square deviation (RMSD) values despite low sequence identity (Figure 2B). With protein sequences of comparable length, this family served as an especially strong test case to identify common principles that differ between *I* and *NI* proteins. Of note, low sequence identity is commonly observed across protein families and underlines the evolutionary capacity of proteins to find alternate sequence solution to the same protein fold (*58*).

**Figure 2:**
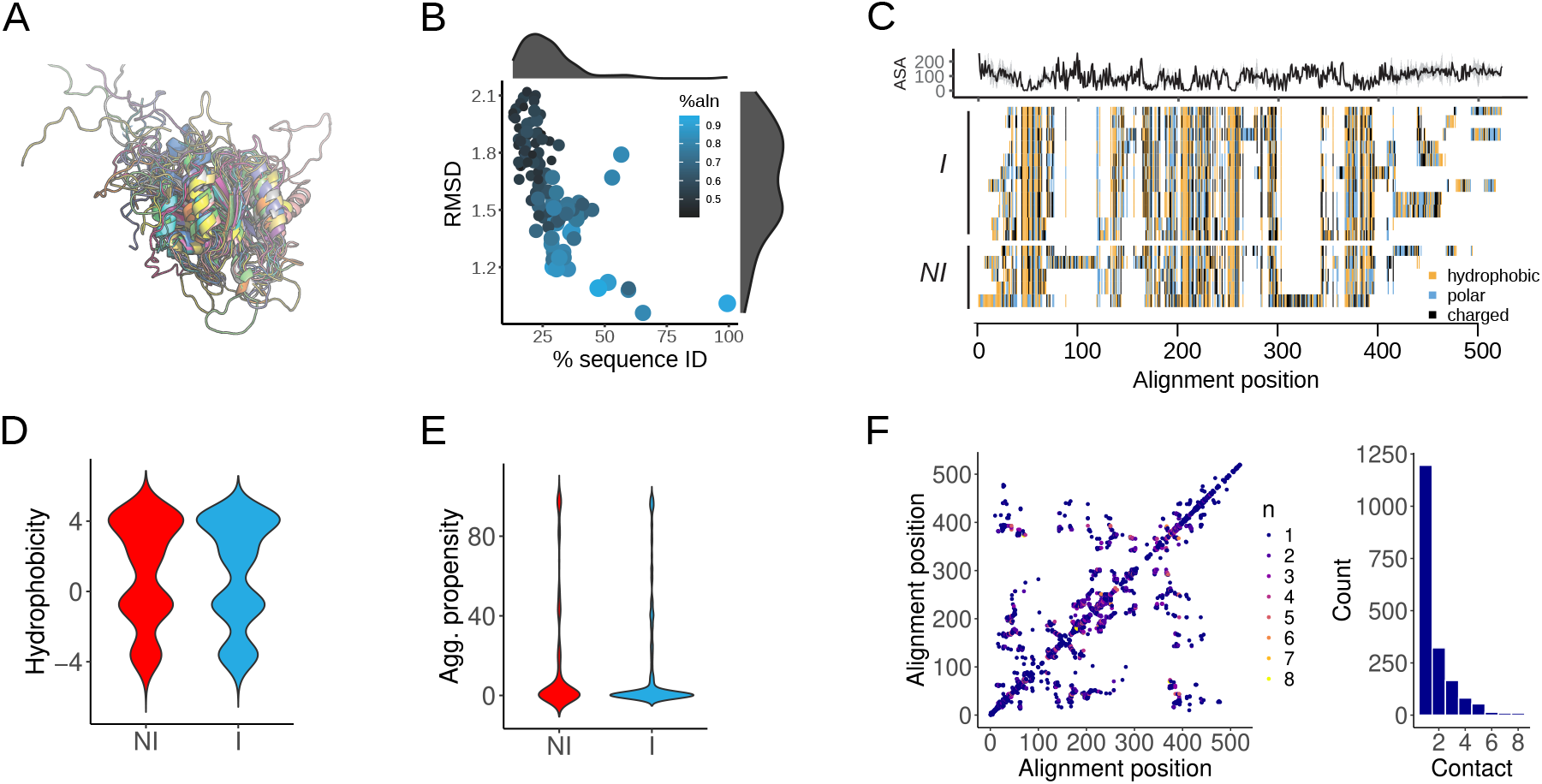
Structural analysis of the G-protein family (c.37.1.8). **A** Superposition of 16 G-protein structures. High structural similarity revealed the conserved structural core. **B** Pairwise RMSD values between G-protein structures as function of their sequence identity and percentage of aligned positions. **C** Visualization of a structure-based multiple sequence alignment for the G-protein family. The average solvent accessible surface area (ASA) identified aligned positions of the structural core. **D** Comparison of sequence hydrophobicity between *I* and *NI* proteins. **E** Comparison of sequence aggregation propensity between *I* and *NI* proteins. **F** Consensus contact map. Hydrophobic contacts in individual structures were projected onto the structurebased multiple sequence alignment and colored according the number of contacts at equivalent positions. A histogram summarizes the counts of contacts that are conserved at equivalent positions.

To understand how these proteins so similar in structure displayed such fundamentally different folding behavior in the cell, i.e. with and without chaperone assistance, first a structurebased multiple sequence alignment of the c.37.1.8 proteins was computed (*see Methods*). Importantly, the obtained alignment clearly identified the sequence regions in individual proteins that constituted the strongly conserved hydrophobic core shared by all proteins (Figure 2C). Low average solvent accessible surface area (ASA) of individual positions directly corresponded to conserved sequence hydrophobicity. In contrast, individual differences between proteins mostly comprised linker regions and surface-exposed loops. However, no immediate difference between *I* and *NI* proteins was apparent.

An increase in protein sequence hydrophobicity or aggregation propensity can increase protein stability but equally render proteins more susceptible to misfolding and aggregation. During *de novo* folding, increased protein sequence hydrophobicity and aggregation propensity initially pose an elevated challenge for folding while avoiding misfolding and aggregation. However, no global differences in sequence hydrophobicity (Figure 2D) or aggregation propensity (Figure 2E) between the c.37.1.8 *I* and *NI* proteins could be observed. The differences in folding behavior thus had to link to different cues. Next, hydrophobic contacts within individual protein structures were mapped onto the structure-based multiple sequence alignment to derive a consensus contact map (Figure 2F, *see Methods*). Remarkably, almost no contacts were conserved across equivalent alignment positions. Evidently, despite the high structural similarity there were fundamental differences in the folding landscapes of these G-proteins (c.37.1.8).

### Contact density links to chaperone interaction

The observation that almost no hydrophobic contacts of the structural core across the c.37.1.8 protein family were conserved opened the possibility that these proteins differed markedly in their folding including their potential for misfolding and formation of non-native contacts. Contact density has long served as a proxy for the complexity of protein folding landscapes (*59*). Buried positions in the structural core were identified based on per-position average ASA (mASA) values of mASA < 50Å^2^, 30Å^2^ and also 20Å^2^ (*60*). The more stringent thresholds reflected that averaging of individual ASA profiles mapped onto the structure-based multiple sequence alignment naturally slightly decreased any values. Notably, at a selective threshold of mASA < 20Å^2^, the contact density in the structural core of NI proteins was found significantly higher than that of the corresponding I proteins (Figure 3A, *p =* 0.027, Wilcoxon-Mann-Whitney (WMW) test). The thresholds of mASA < 30Å^2^ and 50Å^2^ supported the same trend albeit with less pronounced differences and without significance. To explain whether the higher contact density observed in *NI* proteins resulted from a higher number of residues contributing contacts, or from highly connected residues forming more contacts, the degree distributions of core the positions were analyzed. The G-proteins showed a combination of both effects. The *I* proteins had on average both a higher number of core residues that were not forming any contacts, and fewer contact residues that engaged in 1, as well as 2 or more contacts compared to the *NI* proteins (Figure 3C).

**Figure 3:**
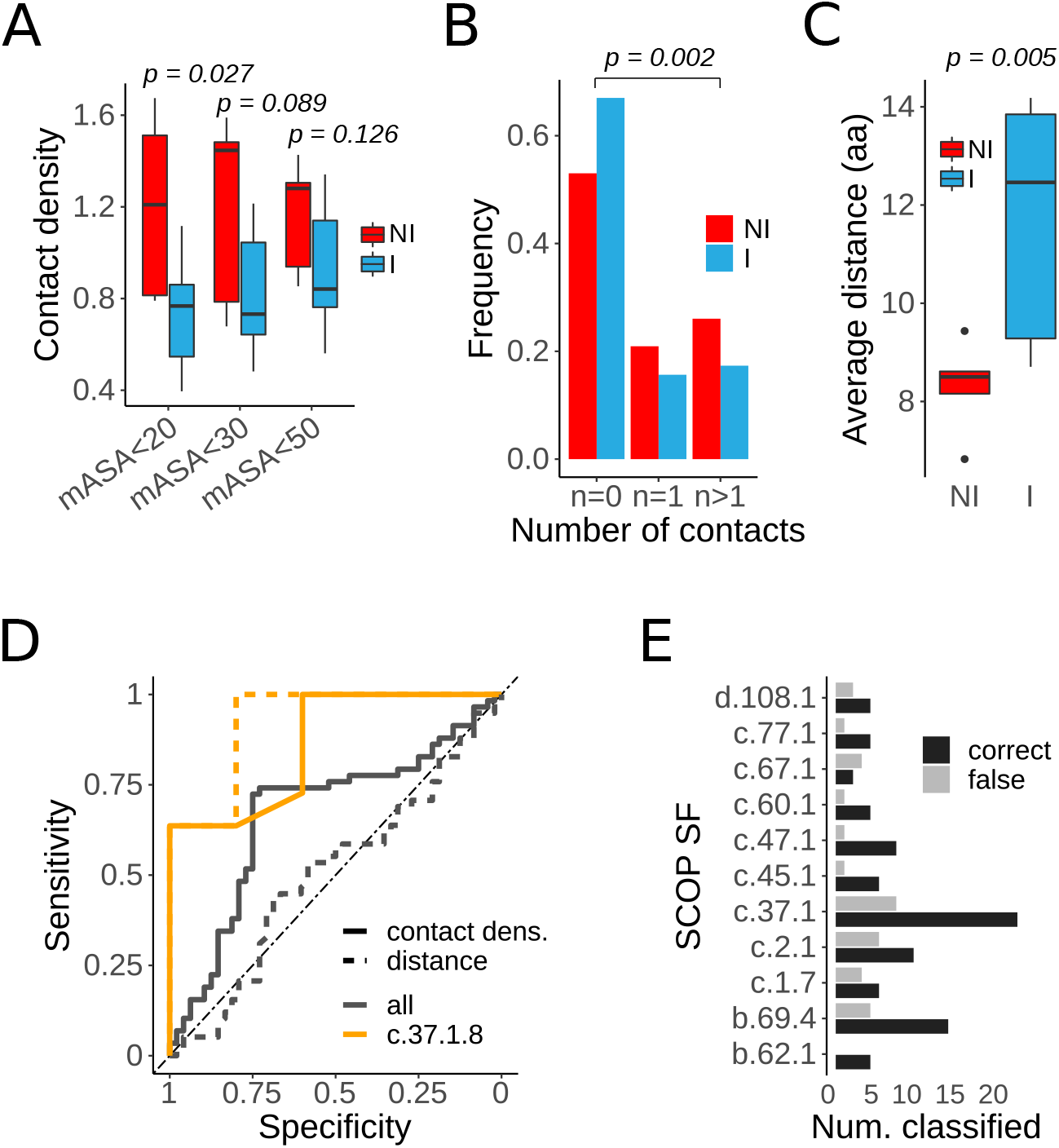
Contact density predicts chaperone interaction. **A** Contact density of buried residues of the structural core of the G-protein family (c.37.1.8). Contact density is based on the number of hydrophobic contacts of positions of average solvent accessible surface area (mASA) values below 20Å^2^, 30Å^2^, and 50Å^2^, respectively. **B** Frequency of residues with no hydrophobic contacts, 1 hydrophobic contact, and 2 or more hydrophobic contacts in the structural core of the G-proteins defined by mASA < 20Å^2^. **C** Distribution of average distances along the protein sequence between buried contacts in the structural core of the G-protein family for *I* and *NI* proteins. **D** Classification of *I* and *NI* proteins based on contact density and contact density. ROC curves for the c.37.1.8 protein family are based on the contact density and contact distance derived from the structure-based multiple sequence alignment. ROC curves for all proteins across superfamilies analyzed are based on contact density and contact distance of buried positions with ASA < 50Å^2^. **E** Overview of correctly and falsely classified proteins by superfamily.

To assess whether an overall lower number of contact residues as well as a higher number of highly connected residues also changed the spacing between sequence positions that formed hydrophobic contacts in the structural core, the average distance between consecutive buried contacts along the protein sequence was computed. Strikingly, the average sequence distance between buried contacts for NI proteins in the c.37.1.8 protein family was significantly shorter than that for I proteins (Figure 3C, p = 0.005, WMW test). Longer distances between contacts can mean a larger conformational search space including a higher possibility of misfolding. Conversely, proteins with shorter distances between buried contacts should be on more ‘funnelled’, or ‘safer’ folding pathways. Intriguingly, the longer distances between buried contacts that could render proteins more susceptible to misfolding matched well the dependency on the Ssb chaperone for de novo folding. Importantly, the observed differences in distances corresponded to length scales relevant for protein folding. The average persistence length in small global proteins has been estimated around 40Å (*61*) or approx. 6 residues. The average distance between contacts in NI proteins thus resembled the persistence length of polypeptide chains, while the average distance between buried contacts in I proteins was clearly longer. Similarly, negative design principles such as strategically placed charges or salt bridges that protect hydrophobic and interaction prone protein segments from aggregation had also been found within average polypeptide persistence length (*54*). Of note, the absolute values of average distances between buried contacts depended on the definition of buried contact residues and were found shorter for less stringent ASA thresholds. Based on these findings, the proteins from the c.37.1.8 family could be accurately classified as quantified based on the area under the curve (AUC) of receiver operating characteristic (ROC) curves and based on their buried contact density (*AUC* = 0.86) and contact distance (*AUC* = 0.93) (Figure 3D).

To test whether this observation was more generally valid, the chaperone interactions across all superfamilies in the assembled data set were predicted based on contact density and distance. Different protein folds impose widely varying constraints on folding. Therefore, contact density values were normalized to be centered around zero as to yield best within superfamily classification. While overall buried contact density maintained discriminative power to predict chaperone interactions, the classification performance for the larger and more disparate data was weaker (Figure 3D, AUC = 0.67). In contract, *I* and *NI* proteins could not be classified based on contact distance across superfamilies (AUC = 0.5). Next to the challenge of comparing proteins from different superfamilies, the identification of buried contacts solely based on ASA values for the full data set as opposed to average ASA values derived from a structurebased multiple sequence alignment may have caused a sufficient loss in resolution. Breaking down the classification of chaperone interaction based on contact density by superfamily, it was noteworthy however that general predictive power could be observed for almost all protein superfamilies (Figure 3F). These results suggested a direct link between the complexity of protein folding landscapes and their dependence on chaperones for de novo folding that is poised to become much more informative through investigation of protein folding dynamics at higher resolution beyond protein contact maps.

### Programmed chaperone interactions

Like any protein interactions, chaperone interactions depend on energetically favorable interactions between complementary sequences or surfaces. The Hsp70 Ssb is known to bind preferentially to short hydrophobic peptide sequences of length 5-7aa (*16,17*). However, well defined recognition motifs could so far not be established (*22*), supported by our own unsuccessful attempts to discover conventional sequence motifs. To test whether Ssb substrates generally contained discriminative sequences that could differentiate between *I* and *NI* proteins, I implemented an algorithm that combined the hierarchical clustering of short linear peptides based on a substitution matrix with Random Forest feature selection to identify clusters of peptide sequences that could predict Ssb interactions (Figure 4A, *see Methods*). Because the identification of discriminative sequences in a small data set may not represent features that determine Ssb interactions proteome wide, the algorithm was first tested on experimentally derived Ssb binding regions in all Ssb interacting sequences (*23*) (*see Methods*). Herein, just over 100,000 unique peptides of 7aa from 1328 binding regions derived from 733 consensus Ssb substrates as well as random fragments from 2393 non-interacting control sequences were clustered and evaluated. Importantly, *I* and *NI* proteins could in this way be clearly classified based on the presence of discriminative short sequences from the most important clusters (Figure 4B,C). Peptide sequences may bind to Ssb in different orientations or interact with slightly different amino acids in the general binding pockets. Therefore, the 1 most important, the 3 most important, and the 5 most important clusters were evaluated representing 1, 3, or 5 different binding modes, respectively. Initial clusters obtained by greedy clustering were small with a lower threshold of 5 sequences. Accordingly, small clusters observed during early hierarchical clustering iterations could not predict well the interaction status of the much larger number of input sequences (Figure 4B). The increasing discriminative power with progressing clustering (Figure 4B) thus initially also linked to growing cluster sizes (Figure 4C). While chaperones including Ssb are thought to bind to hydrophobic sequences, the limiting of the feature selection to only clusters of minimum hydrophobicity (see Methods) lead to a slight drop in classification performance. This result suggested that additional sequence characteristics other than hydrophobicity were important. Towards the end of the hierarchical clustering very large clusters could achieve high classification power (Figure 4B) but at a cost of a clear drop in cluster entropy (Figure 4C). Therefore, these later stage large and heterogeneous clusters of dissimilar peptides likely carry limited to no biological meaning. A closer analysis of the 3 best scoring clusters along the clustering trajectory were very informative. Initial cluster sizes were above average and later below average as to most closely resembling the number of positive and negative sequences of the classification (Figure 4C). Moreover, the best clusters were generally characterized by below average entropy, indicating fundamental limitations in clustering based on a substitution matrix (Figure 4C). Cluster hydrophobicity in turn varied markedly, indicating the possibility of different types of discriminative peptide sequence including directly binding sequences or flanking sequences that confer specificity.

**Figure 4:**
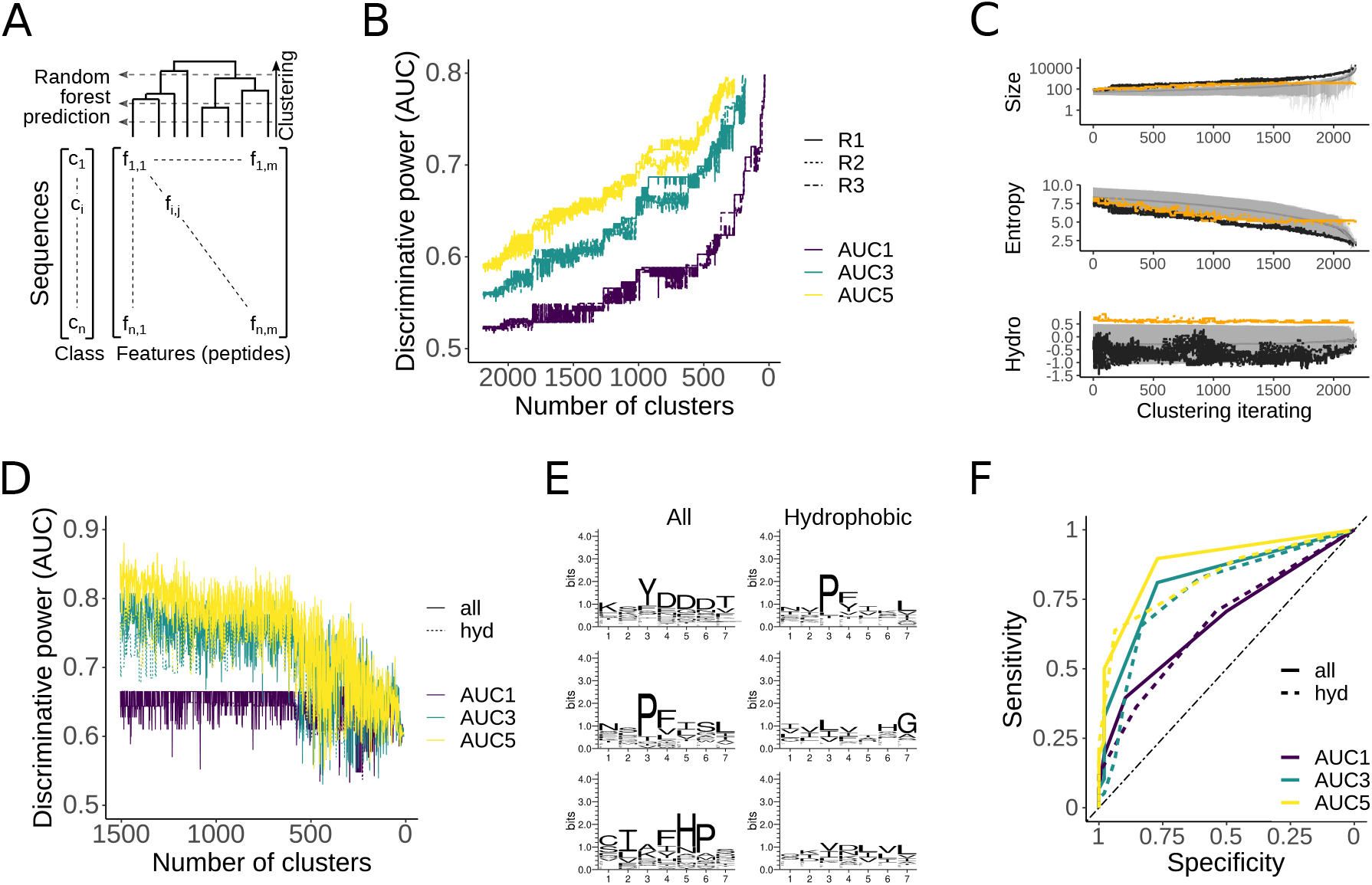
Identification of discriminative peptides through hierarchical clustering and Random Forest feature selection. **A** Schematic of the algorithm. All unique peptide sequences of length 7aa were grouped by hierarchical clustering based on a substitution matrix. At each clustering step, the best clusters of peptides whose presence in protein sequences was predictive of their chaperone interaction were identified by Random Forest feature selection. **B** Identification of discriminative peptides in all yeast Ssb substrates with experimentally determined binding regions. Shown are traces of the best obtained discriminative power at each clustering iteration upon selection the 1, 3, and 5 best clusters, respectively (AUC1, AUC3, AUC5). Because negative control sequences of non-chaperone interactors were based on a randomly sampled subset, three runs (R1, R2, R3) are shown. **C** Summary statistics on the hierarchical clustering of peptide sequences from all Ssb substrates. Averages (grey lines) and standard deviation (shaded area) for cluster size, cluster entropy, and cluster hydrophobicity are shown for three runs as function of clustering iteration. Indicated for reference are the averages of the best 3 performing clusters (black lines), and best 3 performing hydrophobic clusters (orange lines). **D** Identification of discriminative peptides in single-domain Ssb substrates with protein structures. **E** Exemplary sequence logos of best performing clusters of discriminative peptides. **F** Classification of single-domain Ssb substrates based on discriminative sequences. Shown are ROC curves for the best 1, 3, and 5 clusters (AUC1, AUC3, and AUC5) and for considering all clusters or hydrophobic clusters only.

I next sought to test whether discriminative pepdite sequence equally describe Ssb substrates without prior knowledge on the experimentally determined binding regions. The results obtained for the small subset of single-domain Ssb interactors qualitatively matched these observations very well (Figure 4D). The much small data set of approx. 28,000 peptides from 58 *I* and 48 *NI* proteins achieved almost immediate high classification power based on the small clusters from the greedy clustering that subsequently dropped when clusters started becoming too large. The limiting to only hydrophobic clusters resulted in a small drop in performance as observed before, and the broad range of cluster iterations that could achieve comparative classification power suggested that several sequence characteristics were highly characteristic for chaperone interactions. Exemplary sequence logos of best performing clusters range in intermediate information content and represented discriminative sequences that await to be further understood rather than validated binding motifs (Figure 4E). However, the overall classification power based on a small number of peptide sequences derived from unbiased clustering was striking (Figure 4F).

Taken together, this analysis revealed two important insights. Foremost, sparse sequence features had very pronounced discriminative power for the classification of Ssb substrates. This is a fundamental insight that opens the possibility that some chaperone interactions may actually be sequence encoded, i.e. programmed, rather than responding to folding constraints. Moreover, it is well established that peptide sequences often contribute in nonlinear fashion to binding energetics. As a result, the current use of substitution matrices and also entropy metrics are likely insufficient. Instead, peptide sequences should be clustered according to their binding energetics, which is not trivial by may significantly further insights gained from this and related approaches. This notion was further supported by the observation that considering a higher number of clusters readily lead to higher classification power. Thus, any improvement on the similarity between clusters may rapidly improve the overall performance and insights gained.

### Codon usage can compensate for chaperoning

Last, the protein coding mRNA sequences encode an additional layer of information on the kinetics of translation elongation (*62*) thus protein folding (*63, 64*) including interactions with chaperones (*23, 24, 65*). Next to selectively placed signatures of translation attenuation, for instance through use of nonoptimal, more slowly translated codons that can provide extra time for folding events and thus limit the conformational search space, an overall high or low content of nonoptimal codons can bias the overall translation speed. Highly abundant proteins are generally characterized by efficient translation, i.e. enriched in optimal and depleted of nonoptimal codons. Validating any general bias in nonoptimal codon content due to overall protein expression levels, protein abundance data was analyzed (*66*). Normally, chaperone substrates are biased towards long and aggregation prone proteins of low solubility, and as a result expressed at lower levels than non-substrates. Interestingly, the atypical single-domain Ssb interactors displayed opposing behavior. Specifically, I proteins were characterized by significantly higher cellular abundances than their NI counterparts (Figure 5A, *p* = 0.0002, WMW test) that was however not significantly reflected in nonoptimal codon content (Figure 5B, *p* = 0.37, WMW test).

**Figure 5:**
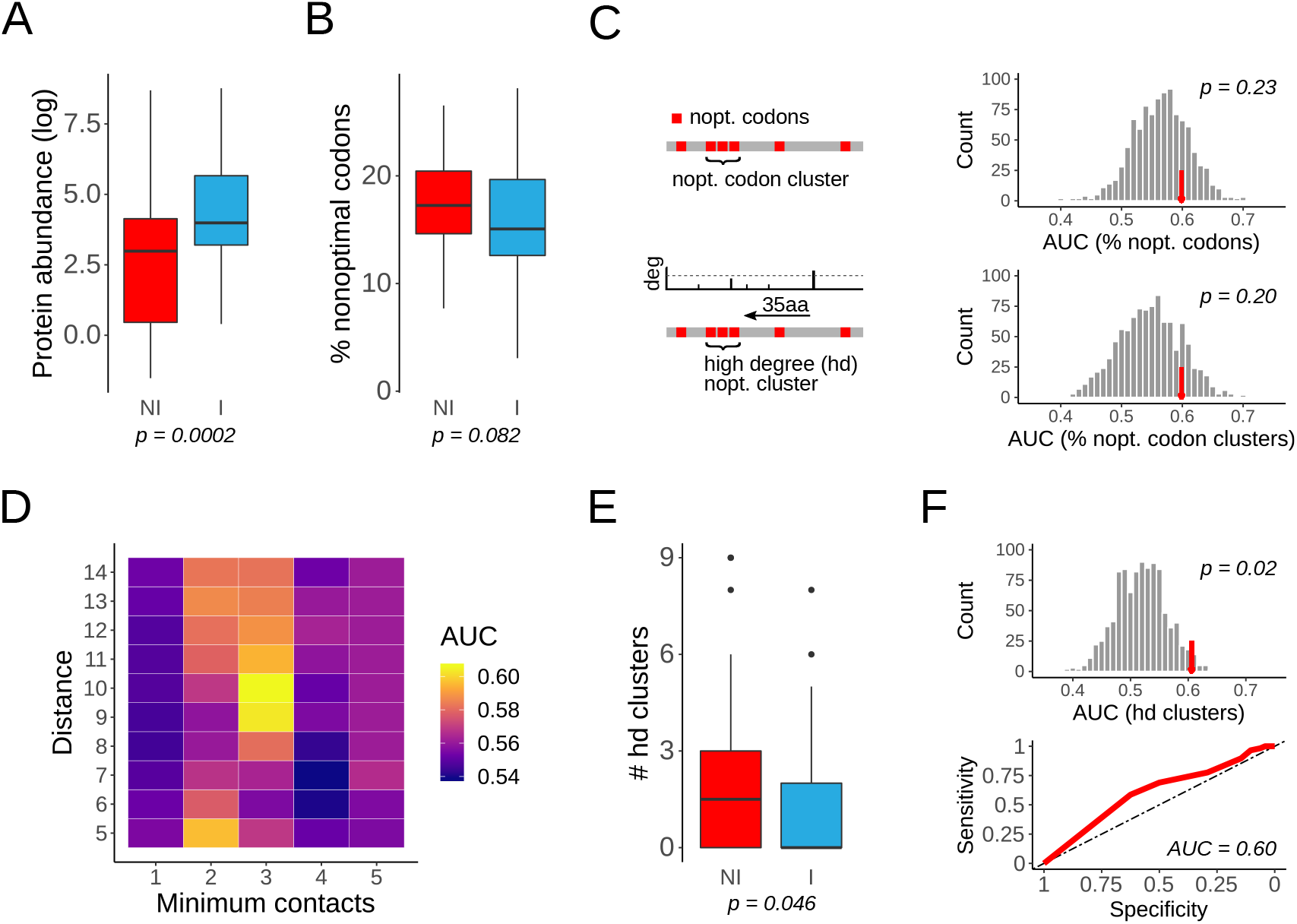
Codon usage can compensate for chaperone interaction. **A** Protein abundances of *I* and *NI* proteins. Contrary to what is normally observed for chaperone substrates, the atypical small single-domain *I* proteins are less expressed than their *NI* counterparts. **B** Nonoptimal codon content. Distributions of the percentages of nonoptimal codons in mRNAs coding for *I* and *NI* proteins. **C** Nonoptimal codon clusters. Quantified are the overall percentage of nonoptimal codons per sequence, the percentage of sequence positions in nonoptimal codon clusters, and per-sequence counts of high-degree (hd) nonoptimal codon clusters found downstream of residues involved in a high number of hydrophobic contacts. The strategic positioning of such nonoptimal codons in clusters may contribute to translational attenuation and thus promote formation of critical hydrophobic contacts outside the ribosome. Both the percentages of nonoptimal codons and the percentages of nonoptimal codon clusters are weakly predictive of Ssb interactions. However, the same discriminative power is readily achieved in synonymous codon randomized sequences, indicating that this observation is not a characteristic of synonymous codon choice. **D** Classification of *I* and *NI* proteins based on the presence of hd nonoptimal codon clusters for different distance and contact thresholds. **E** Distribution of the counts of hd nonoptimal codon clusters in *I* and *NI* proteins. **F** Selection on hd nonoptimal codon clusters. Hd nonoptimal codon clusters in protein sequences are significantly more predictive of Ssb interactions than what can be observed after randomizing synonymous codon usage, suggesting that the positioning of hd nonoptimal codon clusters may be under synonymous codon selection. ROC curve for the classification of *I* and *NI* proteins based on hd nonoptimal codon cluster content indicates weak overall discriminative power.

To better understand if nonoptimal codons was linked to chaperone assistence, different metrics quantifying nonoptimal codon content were evaluated for the discriminative power to predict Ssb interactions. The overall percentage of nonoptimal codons per sequence on its own was weakly predictive of Ssb interactions (*AUC =* 0.60) (Figure 5C). However, upon randomizing synonymous codon usage (see Methods) the similar results could be readily obtained by chance (p = 0.228, empirical p-value), suggesting that the global percentage of nonoptimal codons was not selectively linked to chaperone interaction. Clusters of nonoptimal codons may represent regions in mRNA sequences that are selectively codon-deoptimized. Similarly, the percentage of sequence positions in mRNAs that were part of nonoptimal codon clusters could weakly predict Ssb interactions (AUC = 0.60), although also likely not due to selection on synonymous codon usage (p = 0.197, empirical p-value). While an overall slow translate rate has been linked to elevated protein quality control needs of long and folding challenged proteins (*15,67*), it may be less relevant in the short single-domain proteins analyzed here.

Next, I tested whether there was any link between the positioning of nonoptimal codon clusters and the formation of native protein contacts. The ribosome exit tunnel can protect between 23-50 amino acids of the elongating nascent chain depending on its conformation (*68*). As a result, any analysis of binding or folding events outside the ribosome exit tunnel can only be an average approximation. Thus, accommodating for the length of the ribosome exit tunnel by an average 35 aa, the predictive power of the number of ‘high-degree’ (hd) nonoptimal codon clusters linked to contact formation was tested for different distance and contact thresholds (Figure 5D). While the overall signal was weak and within a narrow range of AUC values between 0.54 and 0.6, the strongest signal could be obtained for hd nonoptimal codon clusters within the tunnel length and an additional 10 amino acids of residues that form 3 or more contacts (Figure 5D). At these parameters, NI proteins had on average a significantly higher content of ‘hd’ nonoptimal codon clusters compared to the I proteins (Figure 5E, p = 0.046, WMW test). Importantly, the discriminative power of the counts of hd nonoptimal codon clusters was lost upon synonymous codon randomization, suggesting that the presence and positioning of hd nonoptimal codon clusters in NI sequence may in part be the direct result of selection on synonymous codon usage (Figure 5E, p = 0.024, empirical p-value). Taken together, the analysis of codon usage suggested only weak predictive power of Ssb interaction in short single-domain proteins but highlighted a potentially important compensatory mechanisms wherein strategically placed nonoptimal codon clusters slow down translation to promote protein folding of hydrophobic contacts in absence of chaperone assistance.

### Trade-offs in protein folding networks

In summary, different complementary contributions to successful protein folding, namely hydrophobic contacts defining the folding landscape, discriminative sequences, and codon usage all contributed to defining Ssb interactions. To quantitatively test whether these different characteristics may directly compensate one and another, Ssb interactions were predicted jointly based on these different features with cross-validated Random Forest models.

Importantly, overall high predictive power and direct coupling between these characteristics could be observed (Figure 6A). When considering all clusters of putative discriminative peptides, the signal was so strong that no gain in classification performance could be achieved upon adding information on contact density and nonoptimal codon clusters. However, when only considering hydrophobic clusters, the overall prediction accuracy further improved by including the contact density and nonoptimal codon cluster features (Figure 6A). This important observation supported the idea that these different features trade-off in compensatory manner. Of note, the high degree nonoptimal codon clusters on their own lost their predictive power upon stringent cross-validation. Similarly, contact density was only weakly predictive upon cross-validation. However, predictions jointly based on contact density and hd nonoptimal codon clusters became very predictive, suggesting very strong alternative and compensatory usage of these features.

**Figure 6:**
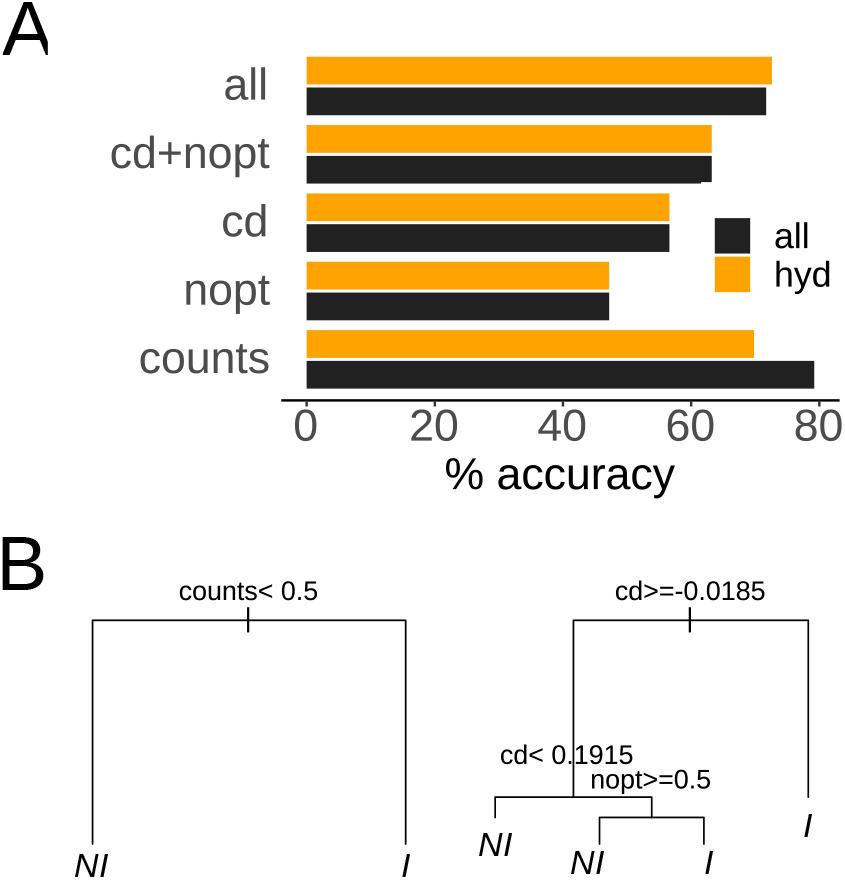
Trade-offs in protein folding networks. **A** Predictive power of Random Forest models upon leave-one-out cross-validation. Contact density (‘cd’), counts of peptides from the 3 most predictive peptide clusters (‘counts’) and counts of high-degre nonoptimal codon clusters (‘nopt’) are used to predict Ssb interactions. Accuracy denotes the percentage of correctly predicted *I* and *NI* proteins. **B** Decision trees summarize trade-offs in protein folding networks. A high number of discriminative peptide sequences as dominant variable for Ssb interaction suggests that some chaperone interactions may be directly sequence encoded rather than responding to folding constraints (left). Omitting the discriminative sequences as classification feature, high contact density can be compensated by strategically placed nonoptimal codon clusters to achieve chaperone independent folding.

Decision trees learned from the feature data further quantified these observation (Figure 6B). Discriminative peptide sequences (‘counts’) readily classified *NI* and *I* proteins. When omitting this dominant variable, direct trade-offs between Ssb interaction and alternative presence of either high contact density (‘cd’) or a high number of high-degree nonoptimal codon clusters (‘nopt’) to compensate for chaperone assistance became immediately visible. These results contribute important insights into the sequence and structure determinants of chaperone interactions, and emphasize the importance of integrated analyses to understand complex biological processes.

## Discussion

Despite their centrality to understanding how cells maintain protein homeostasis, the principles of chaperone specificity and selectivity remain mysterious. Here, a systematic and integrated analysis of the sequence and structural determinants that define the interactions between the S. *cerevisiae* Hsp70 Ssb and 106 single-domain proteins was performed. Chaperone interactions were found to compensate for lower density of hydrophobic contacts within proteins, which may render protein folding landscapes less funnelled and more risky. In turn, clusters of nonoptimal codons strategically placed to slow translation thus promote formation of critical hydrophobic contacts appeared to mitigate the need for chaperone assistance. However, the presence or absence of discriminative peptide sequences was by far the strongest determinant of Ssb interaction. Such peptide sequences may contribute both directly as putative binding sites as well as flanking regions that increase specificity. Importantly, the observation of a strong sequence signal suggested that some chaperone interactions may be directly programmed into the amino acid sequences rather than responding to folding challenges. Next to possibly important functional implications, this finding indicates the potential of incorporating sequence-based constraints of chaperone specificity into whole cell of models protein homeostasis.

Of note, Ssb chaperones preferentially interact with long and folding-challenged multidomain proteins (*15*). The single-domain proteins analyzed here consequently comprised very atypical substrates. This raised the question why some of these small proteins rely on energetically costly chaperones at all given that chaperone independent structural homologues readily existed. In all cases, it was possible that NI proteins interact with chaperones other than Ssb. Especially the topologically complex WD40 repeat-like proteins (b.69.4) likely requires chaperone assistance for folding under many stressful conditions. However, Ssb is by far the most important cotranslationally acting Hsp70 in S. *cerevisiae* (*15,19*). Therefore, the analyzed interaction data should be very representative and, consequently, there must be distinct advantages of chaperone dependency.

An intriguing example of this conundrum is provided by the Poliovirus P1 capsid protein that strictly depends on host cell Hsp90 for folding and capsid assembly (*69*). With its remarkable evolutionary capacity, Poliovirus readily adapts to reduced availability of host cell Hsp90 by an increase in protein stability, reduction in protein aggregation propensity, and selective codon deoptimization to promote folding (*70*). However, P1 is unable to completely loose its chaperone dependence (*69*). While the reasons for this uncompromising dependency are not fully established, it is plausible to speculate about functional advantages. For instance, host cell Hsp90 may serve both as a feedback and feedforward sensor. It’s sequestration by Poliovirus may help the rapid shut down of other host cell functions not important for viral replication. In turn, low or insufficient host cell Hsp90 levels may regulate rebalancing the resource expenditure between host cell viability and viral replication as well as shape evolutionary trajectories. Thus, the dependency on a host cell chaperone likely confers an overall fitness advantage.

While the present work focused on sequence and structure determinants of chaperone interactions, additional pivotal roles are played by the cellular context. The action of several chaperones including Ssb is directly coupled to translation (*71*). Moreover, interaction can depend on localization to specialized ribosome-chaperone complexes (*25*) and ribosome recruitment (*72*). Equally importantly, the mechanisms of action of the chaperones themselves (*73–75*) as well as their coupling to co-chaperones (*76*) can exert further control over their interactions. However, interactions ultimately require favorable energetics between complementary protein sequences and surfaces. Herein, further work is required to rationalize the remarkably broad specificity of chaperones. This includes better understanding complex sequence contributions (*77*) beyond linear sequence motifs (*78*), and the link between protein core, dynamics, and binding specificity (*79, 80*).

The present work also highlights both current opportunity and challenge for structural proteomics and systems biology in general, and for genome-scale models of protein folding networks in particular. The potential benefit of integrating structural data into genome-scale models is undisputed (*81,82*) and rapidly establishing itself as an exciting new frontier of modeling large-scale biochemical networks (*83, 84*). A main challenge in truly bridging the scales from the atomistic to the cellular levels lies in incomplete availability of structural data as well as the development of appropriate structure representations. Proteins are complicated molecules and their behavior often critically depends on tiny details. The strong results on contact density and distance obtained from a high-quality structure-based multiple sequence alignment of a curated protein family immediately dropped in signal when performing the same analysis at lower resolution and more generally only based on structural features (Figure 3). What is currently still missing is a sufficient understanding of sequence and structural context of in vivo folding that would allow direct comparison of heterogeneous sets of proteins based on already available structural data extrapolated and generalized by sequence information. Fortunately, protein folding is ultimately encoded in the genomic sequences. The quest to accelerate structural systems biology is thus directly coupled to better understanding how genomic and protein sequences link to protein folding in the cell.

Last, the multi-faceted dependency of proteins on chaperones encourages to revisit the more general question of why proteins needs chaperones in the first place? Any answer must be at least as varied. Some protein functionality evidently depends on structures or structural assemblies so complex that they can, inside the crowded cellular environment, only be attained with the help of chaperones. Other proteins likely gain different benefits through costly interaction with chaperones such as increased protection or evolutionary plasticity, often reflecting their functional importance within protein networks. Currently underappreciated is a likely central regulatory function of chaperone interactions. As evident from the the dramatic diversification of chaperone families with increasing organism complexity (*13*), individual chaperones with their specific interactions likely play a key role in the selective regulation of cellular networks (*85*). Even seeming similar chaperones have been found with unique roles and distinct specialization given the right context (*86*). This thinking is also supported by the findings that functional innovation through gene duplication often equally rests on increased fidelity of regulation (*89*). Matchingly, the very small and atypical single-domain Ssb substrates analyzed here include a large number of Rho GTPases that contribute important regulatory functions for cell motility (*87, 88*). Validating experiments will be needed to directly establish a regulatory role of their chaperone dependence.

Continued advances in two complementary directions poise for a most exciting outlook. A higher resolution understanding of the link between protein sequences, in vivo folding, and structures will also improve our ability to rationalize the effect of mutations both on individual proteins and on protein networks. And an increase in scope from individual folding networks at the protein level to their collective analysis within cellular protein homeostasis ultimately promises to reveal fundamental insights into cellular regulation, as well as dysregulation in conditions of aging and neurodegeneration.

## Methods and Materials

### Data and computer code availability

Project data and computer code to reproduce all presented results is available at https://github.com/pechmannlab/foldingnets

### Protein structure and interaction data

Two recent data sets on cotranslational interactions of the redundant S. cerevisiae Hsp70 chaperones Ssb1 (*22*) and Ssb2 (*15*) defined putative client proteins as ‘strong’ or ‘weak’ interactors based on their binding enrichment relative to their translation (*15,22*). Here, building a consensus of both data sets, proteins that systematically and strongly interacted with either of Ssb1 or Ssb2 were considered an ‘interactor’ (*I*); proteins that interacted with neither, or only weakly, i.e. sporadically, with at most one of Ssb1 or Ssb2, were considered a ‘non-interactor’ (*NI*). All single-domain *I* and *NI* proteins were grouped based on their SCOP protein superfamily classification (*56, 57*). Proteins from superfamilies with at least two *I* and two *NI* proteins were retained for subsequent analysis, yielding a data set of 110 proteins representing 12 superfamilies.

The structural cordinates of 35 proteins with experimentally determined protein structures were obtained from the Protein Data Bank (PDB) (*90*). For the remaining proteins without experimental protein structures in the PDB a homology model was built with MODELLER (*91*). Using default settings, each protein was modelled 5 times against each experimentally determined protein structure from the same superfamily in the compiled data set as template, if available, else a closely related protein structure from the same superfamily. The overall best model, assessed by the DOPI score (*91*), was selected as the representative structure (Table 1). The e.29.1.2 superfamily with 4 identified single-domain proteins was omitted from further analyses as the proteins varied widely in length, were missing good templates, and consequently no good homology models could be built. The final data set thus included 106 proteins from 11 superfamilies. Missing residues in experimental PDB structures were added by building homology models for the full protein sequences using their native structures as template. Finally, both experimental PDB structures and homology models were checked with the PDBfixer program (*92*) and subjected to a soft molecular dynamics (MD) equilibration in openMM (*93*) with explicit solvent and the Amber force field.

### Structural analysis

Protein secondary structure and solvent accessible surface area (ASA) were assigned with DSSP (*94*). Pairwise similarity between protein structures was assessed through structure superposition and quantified by the *Q* score of the Superpose program (*95*). Protein sequence hydrophobicity was assessed with the Kyte&Doolittle hydrophobicity scale (*96*) normalized to zero mean and unitary standard devation (*54*), and protein sequence aggregation propensity was predicted with Tango (*97*). A structural phylogeny of the SCOP superfamily c.37.1 was constructed from a distance matrix based on the Q scores (*98*). Structure-based multiple sequence alignments were computed with Stamp (*99*). Contact maps of hydrophobic inter-residue contacts were inferred from the PDB structures with the CSU software (*100*). Buried contacts of the structural core of aligned proteins were defined as positions with average ASA values (mASA) below 20Å^2^, 30Å^2^, and 50Å^2^, respectively. Buried contacts in unaligned proteins were identified through ASA values below 50Å^2^. Discriminative power to classify chaperone interactions based on contact density was quantified through the area under the curve (AUC) of the receiver operating characteristic (ROC) analysis with the R package pROC.

### Binding motifs and discriminative sequences

Classical motif finding with the powerful MEME software (*101*) did not yield any significant results that could discriminate between *I* and *NI* proteins, likely because criteria of optimizing motif signal are generally too stringent to describe the broad specificity of chaperones. To simultaneously identify multiple putative binding motifs whose cumulative contribution within sequences can predict their differential interaction with the Hsp70 Ssb, an algorithm that combines the large-scale hierarchical clustering of peptide sequences with Random Forest feature selection was developed. Specifically, positive protein sequences, i.e. sequences known to bind Ssb, and negative sequences, i.e. sequences known to not bind Ssb, were decomposed into all possible short peptide sequences of length *l* = 7 amino acids, the length of peptides bound to the Ssb binding pocket (*16,17*). Each input protein sequence with associated class label of *I* or *NI* was subsequently described by a row of the feature matrix where each column quantified how often a given peptide was found in the protein sequence. Through iterative clustering of peptides in sequence space based on sequence similarity, columns now representing clusters of peptides were derived by summation of the individual peptide counts.

To efficiently cluster large numbers (10^5^ – 10^6^) of short peptides, an initial round of greedy clustering (*102,103*) was performed using the negative of the Blosum62 substitution score as distance between peptides. Starting from the first peptide as initial seed, peptides were assigned to the cluster of the seed peptide with the shortest distance. If no cluster seed within a maximum distance of *d* = —1.7 o l = —11.9 (*103*) could be found, the peptide was established as new and additional cluster seed. The greedy clustering produced small clusters of highly similar peptides while rapidly decreasing the scale of the computational task at hand. All clusters of a minimum size of 5 were retained and further consolidated by standard hierarchical clustering based on average linkage and the negative Blosum62 score as distance metric.

At each hierarchical clustering step, the current peptide clusters were evaluated for their discriminative power to predict the chaperone interaction labels *I* and *NI*. Because chaperone binding sites should be present in *I* proteins and not in *NI* proteins, only clusters whose sum of feature counts across all *I* proteins was greater than that across all *NI* proteins were evaluated. A Random Forest model was fitted and the relative importance of all features (peptide clusters) determined with Python sklearn. To reflect that Hsp70 chaperones may allow binding in different orientations to their binding pockets, the single most important feature, the 3 most important features and the 5 most important features representing 1, 3, and 5 putative binding modes respectively were identified. The discriminative power of the identified features was quantified by the area under the curve (AUC). Because chaperones preferentially bind to hydrophobic sequences, an additional evaluation only considered hydrophobic clusters above an average value of 0.5 based on the Kyte&Doolittle hydrophobicity scale (*96*) normalized to zero mean and unitary standard deviation (*54*).

To better understand how peptide sequences identified in a small subset of proteins may be representative for all Ssb interactions as well as resemble sequences in known Ssb binding regions, the developed clustering algorithm was first tested on the full Ssb interaction data set. Ssb binding regions in proteins were inferred as the regions upstream of mRNA sequences that were translated while Ssb binds to the encoded polypeptide sequence (*23*). Herein, candidate binding regions were considered as the region from the start of the mRNA signal plus 25 positions as lower bound of the sequence inside the ribosome exit tunnel, to the end of the mRNA signal plus 50 positions as upper bound for the ribosome tunnel plus an additional 20 amino acids. A negative control against these candidate binding regions that contain experimentally determined Ssb binding sites was derived from all sequences of NI proteins that did not systematically localize to the endoplasmic reticulum or mitochondria (*104*). For reasons of computational tractability and to balance the data set, random fragments of length 50 amino acids of the negative NI sequences were chosen. While the exact result depends on the randomly chosen control sequences, the obtained results were found qualitatively very consistent (Figure 4). The results were then compared to the clustering of all peptides from the full protein sequences of the data set of this study without prior knowledge on Ssb binding regions.

### Analysis of codon usage

Codons in the lowest 20% of the tRNA adaptation index (tAI) (*105*) were considered as nonoptimal codons. Three characteristics were evaluated for their discriminative power to classify *I* and *NI* proteins, quantified by the AUC of ROC curves: The percentage of nonoptimal codons in the corresponding mRNA sequence as proxy for overall translation efficiency; the percentage of sequence positions occupied by nonoptimal codons in clusters of at least 3 nonoptimal codons within a window of size 4 as proxy of sequence regions that may be selectively codon-deoptimized; and the the number of ‘high degree’ (hd) nonoptimal codon clusters, i.e. clusters of nonoptimal codons positioned downstream from codons that code for amino acids involved in a high number of hydrophobic contacts within the native protein structure. Hd clusters may slow down translation to allow extra time for the formation of critical inter-residue contacts during folding just as hd sites in the nascent polypeptide have emerged from the ribosome exit tunnel. Protein coding mRNA sequences were randomized by assigning synonymous codons with equal probability; to test whether signatures of codon bias may link to selection on synonymous codon usage, 10000 randomizations of all sequences were computed as background model.

## Acknowledgments

This work has been supported through a Discovery grant from the Natural Sciences and Engineering Research Council of Canada, as well as by computing resources managed through Compute Canada and Calcul Québec. SP is grateful to Léonard Sauvé for contributing preliminary results. SP holds the Canada Research Chair in Computational Systems Biology.

